# Polygenic hazard scores in preclinical Alzheimer’s disease

**DOI:** 10.1101/156331

**Authors:** Chin Hong Tan, Leo P. Sugrue, Iris J. Broce, Elizabeth Tong, Jacinth J. X. Tan, Christopher P. Hess, William P. Dillon, Luke W. Bonham, Jennifer S. Yokoyama, Gil D. Rabinovici, Howard J. Rosen, Bruce L. Miller, Bradley T. Hyman, Gerard D. Schellenberg, Lilah M. Besser, Walter A. Kukull, Celeste M. Karch, James B. Brewer, Karolina Kauppi, Linda K. McEvoy, Ole A. Andreassen, Anders M. Dale, Chun Chieh Fan, Rahul S. Desikan

## Abstract

Identifying asymptomatic older individuals at elevated risk for developing Alzheimer’s disease (AD) is of clinical importance. Among 1,081 asymptomatic older adults, a recently validated polygenic hazard score (PHS) significantly predicted time to AD dementia and steeper longitudinal cognitive decline, even after controlling for *APOE* ε4 carrier status. Older individuals in the highest PHS percentiles showed the highest AD incidence rates. PHS predicted longitudinal clinical decline among older individuals with moderate to high CERAD (amyloid) and Braak (tau) scores at autopsy, even among *APOE* ε4 non-carriers. Beyond *APOE*, PHS may help identify asymptomatic individuals at highest risk for developing Alzheimer’s neurodegeneration.

## INTRODUCTION

There is increasing consensus that the pathobiological changes associated with late-onset Alzheimer’s disease (AD) begin years if not decades before the onset of dementia symptoms.^1,2^ Identification of cognitively asymptomatic older adults at elevated risk for AD dementia (i.e. those with preclinical AD) would aid in evaluation of new AD therapies.^2^ Genetic information, such as presence of the ε4 allele of *apolipoprotein E* (*APOE*) can help identify individuals who are at higher risk for AD dementia.^3^ Longitudinal studies have found that *APOE* ε4 status predicts decline to mild cognitive impairment (MCI) and dementia^4^, and steeper cognitive decline in cognitively normal individuals^5^.

Beyond *APOE* ε4 carrier status, recent genetic studies have identified numerous single nucleotide polymorphisms (SNPs), each of which is associated with a small increase in AD dementia risk.^6^ Using genome-wide association (GWAS) from AD cases and controls, we have recently developed a novel ‘polygenic hazard score’ (PHS) for predicting age-specific risk for AD dementia that integrates 31 AD-associated SNPs (including *APOE* ε4) with US-population based AD dementia incidence rates.^7^ To further evaluate the usefulness of the PHS, in this study, we prospectively evaluated whether PHS predicts rate of progression to AD dementia and cognitive decline in cognitively asymptomatic older adults and individuals with MCI.

## METHODS

We evaluated longitudinal clinical and neuropsychological data (from March 2016) from the National Alzheimer’s Coordinating Center (NACC).^8^ Using the NACC uniform dataset, we focused on older individuals classified at baseline as cognitively normal (CN), with a Clinical Dementia Rating^9^ (CDR) of 0 and available genetic information (n =1,081, Table 1). We also evaluated older individuals classified at baseline as MCI (CDR = 0.5) with available genetic information (n = 571, Table 1). We focused on CN and MCI individuals with age of AD dementia onset < age 88 to avoid violations of Cox proportional hazards assumption as evaluated using scaled Schoenfeld residuals (total n = 1,652).

**Table 1.** Cohort demographics

|  | CN (*n* = 1,081) | MCI (*n* = 571) |
| --- | --- | --- |
| Age ± SD | 71.19 (6.65) | 74.70 (5.85) |
| Education ± SD | 16.07 (2.57) | 15.70 (2.91) |
| Sex (% Female) | 719 (66.51) | 291 (50.96) |
| *APOE* ε4 carriers (%) | 297 (27.47) | 347 (60.77) |
| Converted to AD dementia (%) | 38 (3.52) | 390 (68.30) |
| Baseline MMSE ± SD | 29.22 (1.05) | 25.67 (3.36) |

*MMSE: Mini-Mental State Examination*

We first investigated the effects of the PHS on progression to AD dementia by using a Cox proportional hazards model, with time to event indicated by age of AD dementia onset. We resolved ‘ties’ using the Breslow method. We co-varied for effects of sex, *APOE* ε4 status (binarized as having at least 1 ε4 allele versus none), education and age at baseline. To prevent violating the proportional hazards assumptions, we additionally stratified baseline age by quintiles.^10^

Next, we employed a linear mixed-effects (LME) model to evaluate the relationship between PHS and longitudinal clinical decline as assessed by change in CDR-Sum of Boxes (CDR-SB) as well as by change in Logical Memory test (LMT), Wechsler Adult Intelligence Scale-Revised (WAIS-R) Digit Symbol, the Boston Naming Test (BNT), Trail-Making Tests A and B (TMTA/B), forward and backward Digit Span (f/b DST) tests. To maintain consistent directionality across all tests, we inverted the scale for Trail-Making tests such that lower scores represent decline. We co-varied for sex, *APOE* ε4 status, education, baseline age and all their respective interactions with time. We examined the main effect of PHS by comparing slopes of cognitive decline over time in the neuropsychological tests for individuals with high (~84 percentile) and low PHS (~16 percentile), defined by 1 standard deviation above or below the mean of PHS respectively.^11^ We also compared goodness of fit between the LME models with and without PHS using likelihood ratio tests.

Finally, we evaluated the relationship between PHS, *APOE* and neuropathology in preclinical AD. Specifically, we conducted LME analysis assessing longitudinal change in CDR-SB in CN individuals with available neuropathology (specifically, neuritic plaque scores based on the Consortium to Establish a Registry for AD (CERAD) and neurofibrillary tangle scores assessed with Braak stages).

## RESULTS

PHS significantly predicted risk of progression from CN to AD dementia (hazard ratio (HR) = 2.36, 95% confidence interval (CI) = 1.38 – 4.03, *p* = 1.66×10^-3^) illustrating that polygenic information beyond *APOE* ε4 can identify asymptomatic older individuals at greatest risk for developing AD dementia. Individuals in the highest PHS decile had the highest annualized AD incidence rates (Figure 1). PHS also significantly influenced risk of progression to AD dementia in MCI individuals (HR = 1.17, 95% CI = 1.02 – 1.35, *p* = 2.36×10^-2^). Using the combined CN and MCI cohorts (total n = 1,652) to maximize statistical power, we found that PHS significantly predicted risk of progression from CN and MCI to AD dementia (HR = 1.31, 95% CI = 1.14 – 1.51, *p* = 1.82×10^-4^) (Supplemental Figure 1). At 50% risk for progressing to AD dementia, the expected age for developing AD dementia is approximately 85 years for an individual with low PHS (~16 percentile); however, for an individual with high PHS (~84 percentile), the expected age of onset is approximately 78 years (Supplemental Figure 1). In all Cox models, the proportional hazard assumption was valid for all covariates.

**Figure 1.**
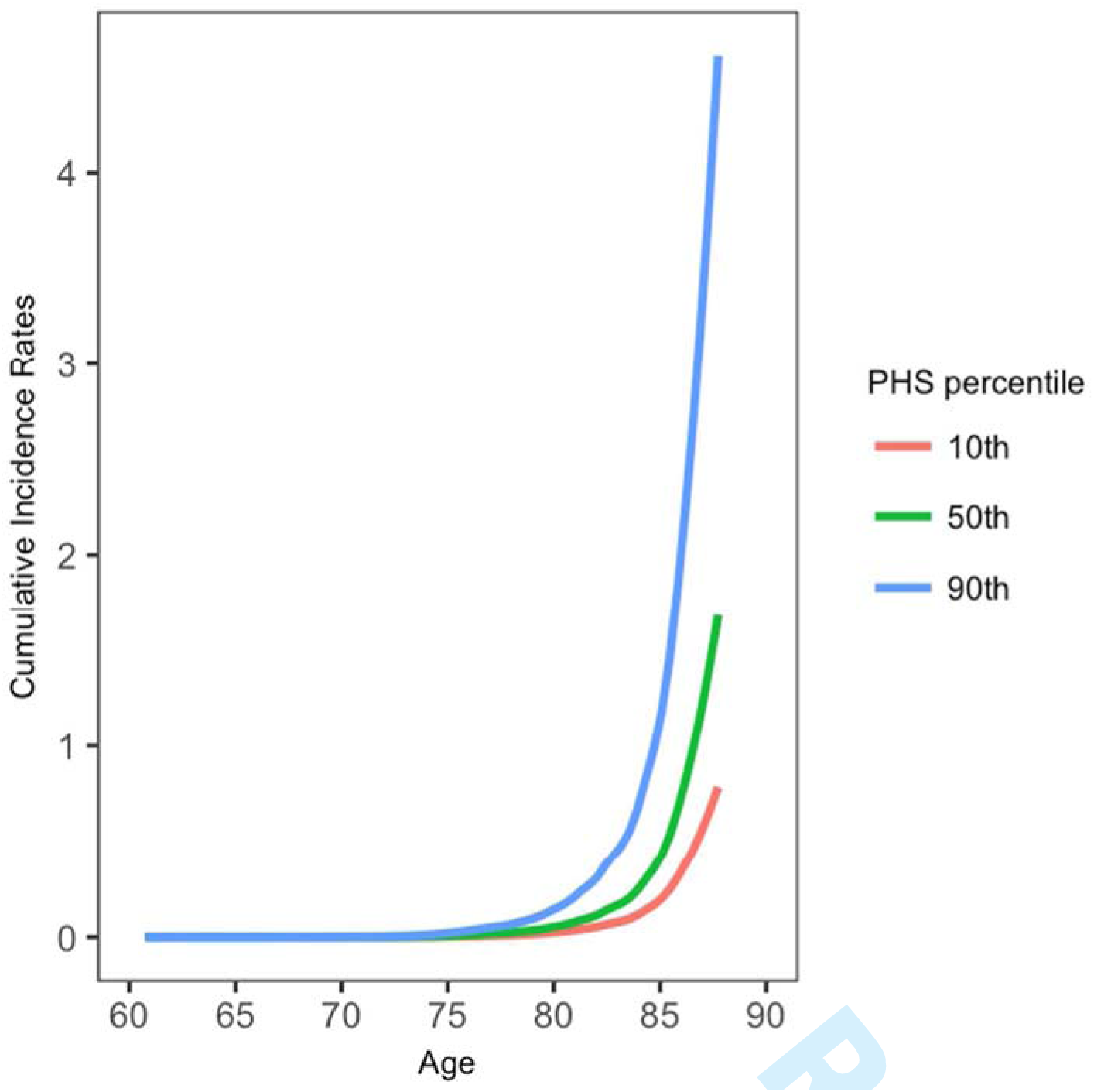
Annualized or cumulative incidence rates in CN individuals showing the instantaneous hazard as a function of PHS percentiles and age.

Evaluating clinical progression and cognitive decline within the CN individuals, we found significant PHS by time interactions for CDR-SB (ß = 0.05, standard error (SE) = 0.02, p = 3.64×10^-4^), WAIS-R (ß = -0.61, SE = 0.30, p = 4.25×10^-2^), TMTB (ß = -2.48, SE = 0.99, p = 1.20 ×10^-2^), and fDST test (ß = -0.93, SE = 0.45, p = 3.76×10^-2^) (Supplemental Table 1), with significantly steeper slopes for high PHS individuals for WAIS-R, TMTB, and CDR-SB (Supplemental Table 2, Figure 2). Evaluating average percentage change across all neuropsychological tests, we found that PHS predicted cognitive decline (ß = 0.84, SE = 0.30, *p* = 4.50×10^-3^), with high PHS individuals showing greater rates of decline (ß = -1.80, SE = 0.89, *p* = 4.30×10^-2^) compared to low PHS individuals (ß = -0.12, SE = 0.80, *p* = 0.88). Goodness of fit comparison using likelihood ratio tests showed that the full LME model comprising PHS and covariates resulted in a better model fit for predicting decline in CDR-SB, BNT, WAIS-R, fDST and TMTB (Supplemental Table 3). We found similar results within the MCI individuals and the combined CN and MCI cohort (Supplemental Tables 1-7, Supplemental Figure 2) illustrating that polygenic information beyond *APOE* ε4 can identify asymptomatic and mildly symptomatic individuals at highest risk for clinical and cognitive decline.

**Figure 2.**
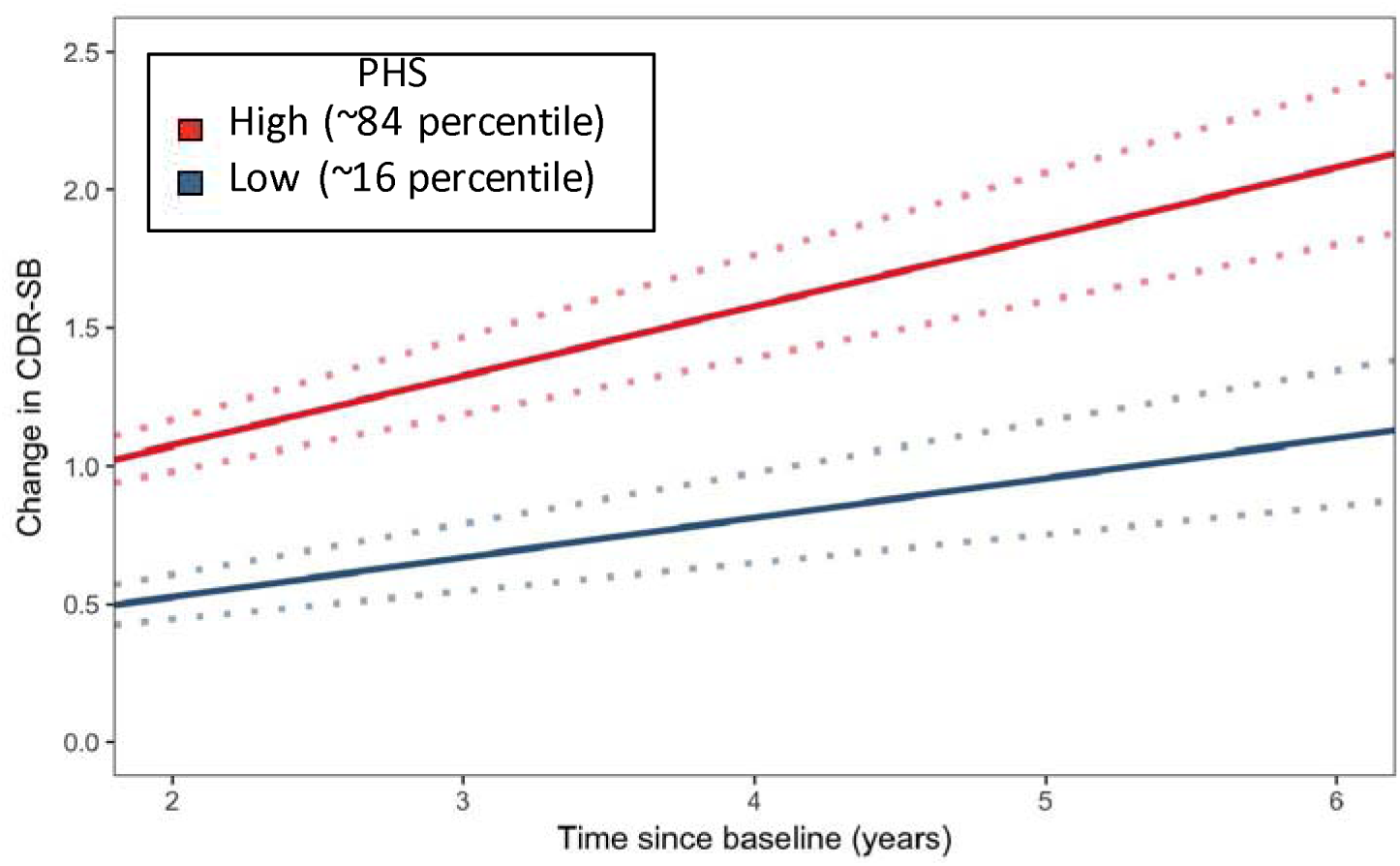
Differences in change over time in CDR-SB in CN individuals over time for low (-1 SD, ~16 percentile) and high (+1 SD, ~84 percentile) polygenic hazard score (PHS) individuals. Dotted lines around fitted line indicate estimated standard error.

Finally, among CN individuals with moderate and frequent CERAD “C” score at autopsy, we found that PHS predicted change in CDR-SB over time (ß = 1.25, SE = 0.28, *p* = 6.63 ×10^-6^), with high PHS individuals showing a greater rate of increase (ß = 5.62, SE = 0. 92, *p* = 1.23×10^-9^). In a reduced model without PHS, *APOE* ε4 status did not predict change in CDR-SB (ß = 0.26, SE = 0.50, *p* = 0.61). Furthermore, even in *APOE* ε4 non-carriers, PHS predicted change in CDR-SB over time (ß = 2.11, SE = 0.38, *p* = 3.06×10^-8^), with high PHS individuals showing a greater rate of increase (ß = 6.11, SE = 1.08, *p* = 1.60×10^-8^). Similarly, among CN individuals with Braak stage III – IV at autopsy, PHS predicted change in CDR-SB over time (ß = 0.93, SE = 0.24, *p* = 1.11 ×10^-4^), with high PHS individuals showing a greater rate of increase (ß = 3.98, SE = 0.79, *p* = 4.45×10^-7^).

**DISCUSSION**

Here, we show that PHS predicts time to progress to AD dementia and longitudinal cognitive decline in both preclinical AD and MCI. Among CN individuals with moderate to high CERAD and Braak scores at autopsy, we found that PHS predicted longitudinal clinical decline, even among *APOE* ε4 non-carriers. Beyond *APOE*, our findings indicate that PHS can be useful to identify asymptomatic older individuals at greatest risk for developing AD neurodegeneration.

These results illustrate the value of leveraging the polygenic architecture of the Alzheimer’s disease process. Building on prior work^4,5^, our findings indicate that polygenic information may be more informative than *APOE* for predicting clinical and cognitive progression in preclinical AD. Although prior studies have used polygenic risk scores in preclinical AD,^12^^-^^14^ by focusing on maximizing differences between ‘cases’ and ‘controls’, this approach is clinically suboptimal for assessing an age dependent process like AD dementia where a subset of ‘controls’ will develop dementia over time (see Figure 1). Furthermore, given the bias for selecting diseased cases and normal controls, baseline hazard/risk estimates derived from GWAS samples cannot be applied to older individuals from the general population. ^15^ By employing an age-dependent, survival analysis framework and integrating AD-associated SNPs with established population-based incidence rates^16^ PHS provides an accurate estimate of age of onset risk in preclinical AD.

In our neuropathology analyses, PHS predicted longitudinal clinical decline in older individuals with moderate to high amyloid or tau pathology indicating that PHS may serve as an enrichment strategy for secondary prevention trials. Congruent with recent findings that the risk of dementia among *APOE* ε4/4 is lower than previously estimated^17^, among CNs with moderate to high amyloid load, we found that *APOE* did not predict clinical decline and PHS predicted change in CDR-SB even among *APOE* ε4 non-carriers. Our combined findings suggest that beyond *APOE*, PHS may prove useful both as a risk stratification and enrichment marker to identify asymptomatic individuals most likely to develop Alzheimer’s neurodegeneration.

## ACKNOWLEDGEMENTS

We thank the Shiley-Marcos Alzheimer’s Disease Research Center at UCSD, UCSF Memory and Aging Center and UCSF Center for Precision Neuroimaging for continued support. This work was supported by the RSNA Resident/Fellow Award, ASNR Foundation AD Imaging Award, NACC JI award, National Institutes of Health grants (NIH-AG046374, K01AG049152), the Research Council of Norway (#213837, #225989, #223273, #237250/EU JPND), the South East Norway Health Authority (2013-123), Norwegian Health Association and the KG Jebsen Foundation. Please see Supplemental Acknowledgements for NIAGADS, ADGC, and NACC funding sources.

**Conflicts of Interest Disclosures:** JBB served on advisory boards for Elan, Bristol-Myers Squibb, Avanir, Novartis, Genentech, and Eli Lilly and holds stock options in CorTechs Labs, Inc. and Human Longevity, Inc. LKM holds stock in CorTechs Labs, Inc. AMD is a founder of and holds equity in CorTechs Labs, Inc., and serves on its Scientific Advisory Board. He is also a member of the Scientific Advisory Board of Human Longevity, Inc. (HLI), and receives research funding from General Electric Healthcare (GEHC). The terms of these arrangements have been reviewed and approved by the University of California, San Diego in accordance with its conflict of interest policies.

## SUPPLEMENTAL FIGURE LEGENDS

**Supplemental Figure 1.**
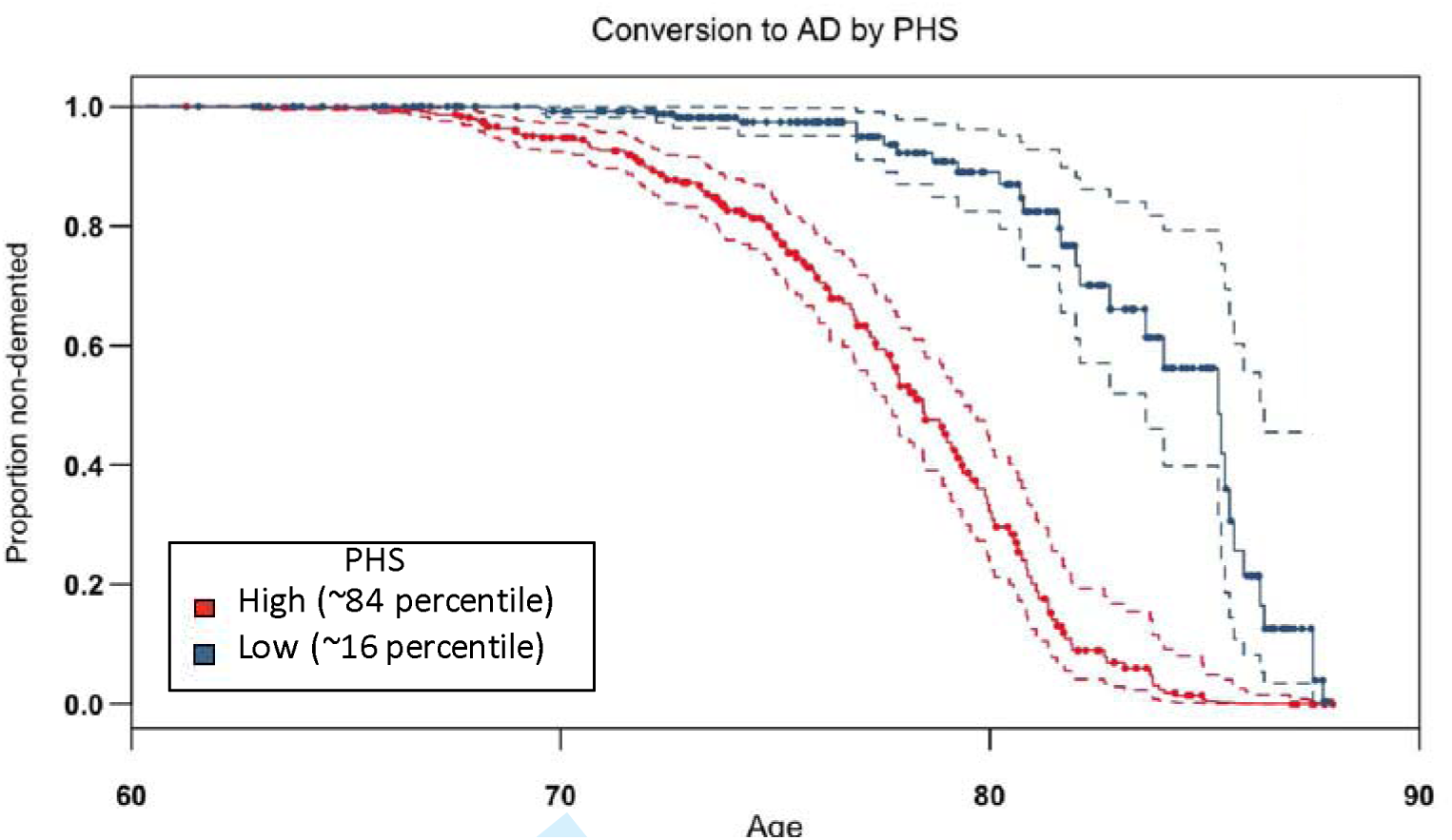
Survivor plot showing progression to AD dementia for low (-1 SD, ~16 percentile) and high (+1 SD, ~84 percentile) polygenic hazard score (PHS) individuals who were CN and MCI at baseline. Dotted lines represent 95% confidence intervals.

**Supplemental Figure 2.**
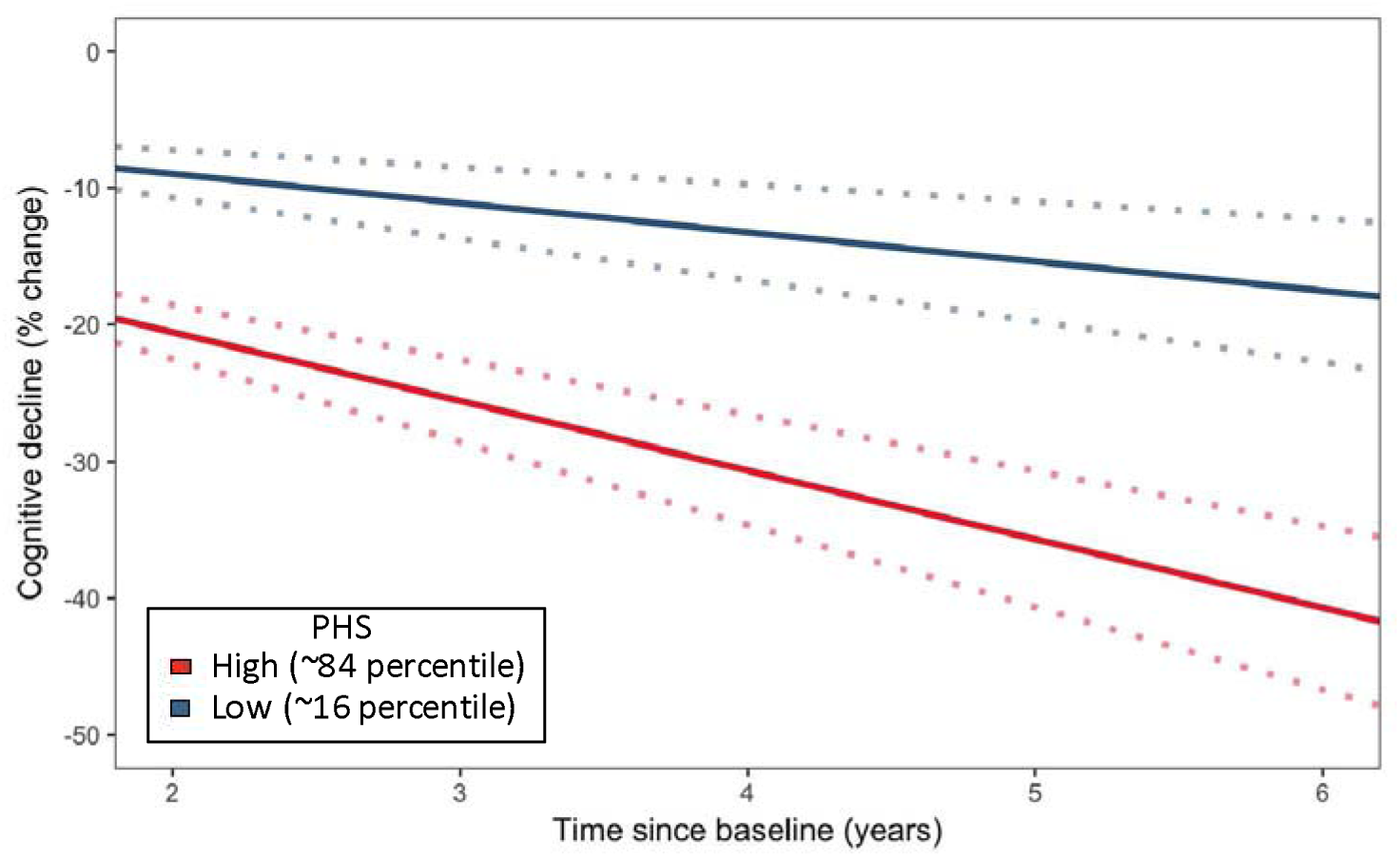
Differences in percentage change in cognitive performance in the combined CN + MCI cohort (average over all neuropsychological tests) over time for low (-1 SD, ~16 percentile) and high (+1 SD, ~84 percentile) polygenic hazard score (PHS) individuals. Dotted lines around fitted line indicate estimated standard errors.

**Supplementary Table 1.**
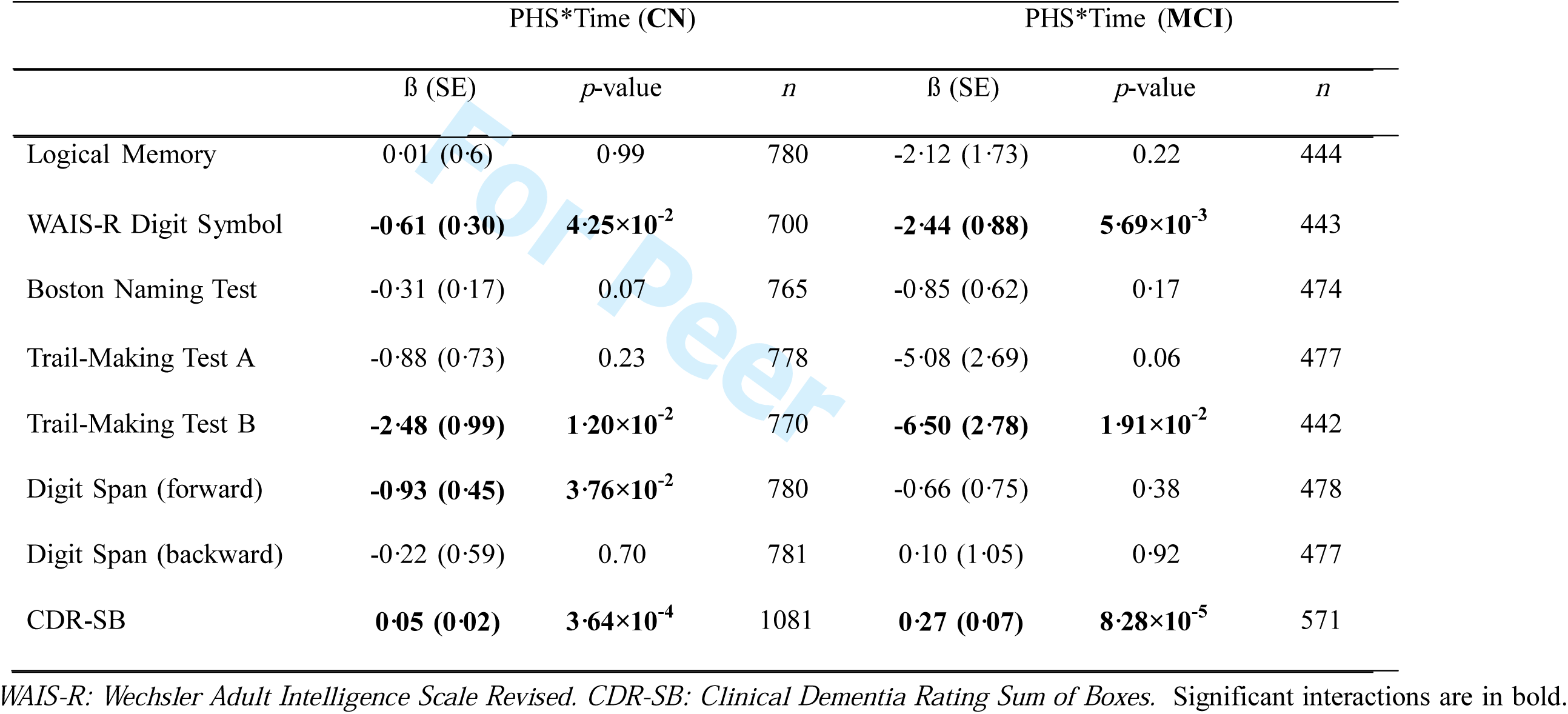
Effects of polygenic hazard score (PHS) on longitudinal cognitive decline in CN and MCI individuals separately.

**Supplementary Table 2.** Differences in rates of cognitive decline for low and high polygenic hazard score (PHS) in CN individuals.

|  | Low PHS (~16 percentile) |  | High PHS (~84 percentile) |  |
| --- | --- | --- | --- | --- |
| | $\beta$ (SE) | *p*-value | $\beta$ (SE) | *p*-value |
| Logical Memory | 1.97 (1.61) | 0.22 | 1.99 (1.82) | 0.27 |
| WAIS-R Digit Symbol | **-2.08 (0.82)** | **1.08×10<sup>-2</sup>** | **-3.30 (0.90)** | **2.66×10<sup>-4</sup>** |
| Boston Naming Test | -0.18 (0.46) | 0.69 | -0.81 (0.51) | 0.11 |
| Trail-Making Test A | -2.19 (1.95) | 0.26 | -3.96 (2.19) | 0.07 |
| Trail-Making Test B | -0.85 (2.62) | 0.75 | **-5.81 (2.96)** | **4.94×10<sup>-2</sup>** |
| Digit Span (forward) | 0.03 (1.19) | 0.98 | -1.82 (1.33) | 0.17 |
| Digit Span (backward) | -1.13 (1.57) | 0.47 | -1.58 (1.77) | 0.37 |
| CDR-CB | **0.14 (0.04)** | **4.28×10<sup>-4</sup>** | **0.25 (0.05)** | **<6.03×10<sup>-8</sup>** |

*WAIS-R: Wechsler Adult Intelligence Scale Revised. CDR-SB: Clinical Dementia*
*Rating Sum of Boxes.* Significant interactions are in bold.

**Supplementary Table 3.** Goodness of fit improvements for linear mixed-effects models with the addition of polygenic hazard score (PHS) in CN and MCI individuals separately using likelihood ratio tests.

|  | CN |  | MCI |  |
| --- | --- | --- | --- | --- |
| | $\chi^2(2)$ | *p*-value | $\chi^2(2)$ | *p*-value |
| Logical Memory | 0.01 | 0.99 | 2.11 | 0.35 |
| WAIS-R Digit Symbol | **7.08** | **2.91×10<sup>-2</sup>** | **10.84** | **4.43×10<sup>-4</sup>** |
| Boston Naming Test | **8.78** | **1.24×10<sup>-2</sup>** | 2.33 | 0.31 |
| Trail-Making Test A | 4.01 | 0.13 | **7.20** | **2.73×10<sup>-2</sup>** |
| Trail-Making Test B | **14.42** | **7.40×10<sup>-4</sup>** | **8.84** | **1.20×10<sup>-2</sup>** |
| Digit Span (forward) | **6.97** | **3.07×10<sup>-2</sup>** | 0.95 | 0.62 |
| Digit Span (backward) | 0.17 | 0.92 | 1.00 | 0.61 |
| CDR-SB | **29.30** | **4.34×10<sup>-7</sup>** | **22.47** | **1.32×10<sup>-5</sup>** |

*WAIS-R: Wechsler Adult Intelligence Scale Revised. CDR-SB: Clinical Dementia Rating Sum of Boxes.* Significant interactions are in bold.

**Supplementary Table 4.** Differences in rates of cognitive decline for low and high polygenic hazard score (PHS) in MCI individuals.

|  | Low PHS (~16 percentile) |  | High PHS (~84 percentile) |  |
| --- | --- | --- | --- | --- |
| | $\beta$ (SE) | *p*-value | $\beta$ (SE) | *p*-value |
| Logical Memory | -7.74 (4.30) | 0.07 | **-11.99 (5.32)** | **$2.44 \times 10^{-2}$** |
| WAIS-R Digit Symbol | **-9.10 (2.19)** | **$3.19 \times 10^{-5}$** | **-13.97 (2.70)** | **$2.29 \times 10^{-7}$** |
| Boston Naming Test | **-5.58 (1.50)** | **$2.04 \times 10^{-4}$** | **-7.28 (1.88)** | **$1.11 \times 10^{-4}$** |
| Trail-Making Test A | **-21.48 (6.50)** | **$9.57 \times 10^{-4}$** | **-31.64 (8.14)** | **$1.01 \times 10^{-4}$** |
| Trail-Making Test B | -11.76 (6.29) | 0.06 | **-24.78 (8.03)** | **$2.02 \times 10^{-3}$** |
| Digit Span (forward) | -2.71 (1.85) | 0.14 | -4.04 (2.28) | 0.08 |
| Digit Span (backward) | **-6.49 (2.58)** | **$1.19 \times 10^{-2}$** | **-6.27 (3.18)** | **$4.89 \times 10^{-3}$** |
| CDR-CB | **0.72 (0.18)** | **$8.89 \times 10^{-5}$** | **1.27 (0.22)** | **$1.13 \times 10^{-8}$** |

WAIS-R: Wechsler Adult Intelligence Scale Revised. CDR-SB: Clinical Dementia
Rating Sum of Boxes. Significant interactions are in bold.

**Supplementary Table 5.** Effects of polygenic hazard score (PHS) on longitudinal cognitive decline in CN and MCI individuals combined.

|  | PHS*Time |  |  |
| --- | --- | --- | --- |
| | $\beta$ (SE) | *p*-value | *n* |
| Logical Memory | **-1.51 (0.68)** | **$2.74 \times 10^{-2}$** | 1,224 |
| WAIS-R Digit Symbol | **-1.53 (0.35)** | **$1.60 \times 10^{-5}$** | 1,143 |
| Boston Naming Test | **-0.96 (0.24)** | **$6.98 \times 10^{-5}$** | 1,239 |
| Trail-Making Test A | **-3.25 (1.00)** | **$1.05 \times 10^{-3}$** | 1,255 |
| Trail-Making Test B | **-4.36 (1.04)** | **$2.85 \times 10^{-5}$** | 1,212 |
| Digit Span (forward) | **-1.07 (0.39)** | **$5.47 \times 10^{-3}$** | 1,258 |
| Digit Span (backward) | -0.59 (0.53) | 0.26 | 1,258 |
| CDR-SB | **0.22 (0.03)** | **$8.88 \times 10^{-16}$** | 1652 |

*WAIS-R: Wechsler Adult Intelligence Scale Revised. CDR-SB: Clinical Dementia Rating Sum of Boxes. Significant interactions are in bold.*

**Supplementary Table 6.** Differences in rates of cognitive decline for low and high polygenic hazard score (PHS) in CN and MCI individuals combined.

|  | Low PHS (~16 percentile) |  | High PHS (~84 percentile) |  |
| --- | --- | --- | --- | --- |
| | $\beta$ (SE) | *p*-value | $\beta$ (SE) | *p*-value |
| Logical Memory | -3.20 (1.80) | 0.07 | **-6.22 (2.06)** | **$2.54 \times 10^{-3}$** |
| WAIS-R Digit Symbol | **-5.38 (0.94)** | **$1.20 \times 10^{-8}$** | **-8.44 (1.07)** | **$2.89 \times 10^{-15}$** |
| Boston Naming Test | **-3.02 (0.62)** | **$1.28 \times 10^{-6}$** | **-4.94 (0.72)** | **$5.01 \times 10^{-12}$** |
| Trail-Making Test A | **-10.95 (2.58)** | **$2.12 \times 10^{-5}$** | **-17.46 (2.96)** | **$3.79 \times 10^{-9}$** |
| Trail-Making Test B | **-6.08 (2.67)** | **$2.26 \times 10^{-2}$** | **-14.81 (3.07)** | **$1.46 \times 10^{-6}$** |
| Digit Span (forward) | -1.28 (1.01) | 0.21 | **-3.43 (1.16)** | **$3.05 \times 10^{-3}$** |
| Digit Span (backward) | **-3.90 (1.38)** | **$4.54 \times 10^{-3}$** | **-5.09 (1.56)** | **$1.27 \times 10^{-3}$** |
| CDR-CB | **0.63 (0.07)** | **$<2 \times 10^{-16}$** | **1.08 (0.09)** | **$<2 \times 10^{-16}$** |

*WAIS-R: Wechsler Adult Intelligence Scale Revised. CDR-SB: Clinical Dementia*
*Rating Sum of Boxes.* Significant interactions are in bold.

**Supplementary Table 7.** Goodness of fit improvements for linear mixed-effects models with the addition of polygenic hazard score (PHS) in CN and MCI individuals combined using likelihood ratio tests.

| | $\chi^2(2)$ | *p*-value |
| --- | --- | --- |
| Logical Memory | **6·85** | **3·26×10<sup>-2</sup>** |
| WAIS-R Digit Symbol | **28·16** | **7·67×10<sup>-7</sup>** |
| Boston Naming Test | **21·74** | **1·91×10<sup>-5</sup>** |
| Trail-Making Test A | **21·58** | **2·06×10<sup>-5</sup>** |
| Trail-Making Test B | **34·41** | **3·38×10<sup>-7</sup>** |
| Digit Span (forward) | **10·78** | **4·57×10<sup>-3</sup>** |
| Digit Span (backward) | 2·78 | 0·25 |
| CDR-SB | **88·33** | **<2×10<sup>-16</sup>** |

*WAIS-R: Wechsler Adult Intelligence Scale Revised. CDR-SB: Clinical Dementia Rating Sum of Boxes.* Significant interactions are in bold.

## SUPPLEMENTAL ACKNOWLEDGEMENTS

**ADGC:** The National Institutes of Health, National Institute on Aging (NIH-NIA) supported this work through the following grants: ADGC, U01 AG032984, RC2 AG036528; NACC, U01 AG016976; NCRAD, U24 AG021886; NIA LOAD, U24 AG026395, U24 AG026390; Banner Sun Health Research Institute P30 AG019610; Boston University, P30 AG013846, U01 AG10483, R01 CA129769, R01 MH080295, R01 AG017173, R01 AG025259, R01AG33193; Columbia University, P50 AG008702, R37 AG015473; Duke University, P30 AG028377, AG05128; Emory University, AG025688; Group Health Research Institute, UO1 AG06781, UO1 HG004610; Indiana University, P30 AG10133; Johns Hopkins University, P50 AG005146, R01 AG020688; Massachusetts General Hospital, P50 AG005134; Mayo Clinic, P50 AG016574; Mount Sinai School of Medicine, P50 AG005138, P01 AG002219; New York University, P30 AG08051, MO1RR00096, UL1 RR029893, 5R01AG012101, 5R01AG022374, 5R01AG013616, 1RC2AG036502, 1R01AG035137; Northwestern University, P30 AG013854; Oregon Health & Science University, P30 AG008017, R01 AG026916; Rush University, P30 AG010161, R01 AG019085, R01 AG15819, R01 AG17917, R01 AG30146; TGen, R01 NS059873; University of Alabama at Birmingham, P50 AG016582, UL1RR02777; University of Arizona, R01 AG031581; University of California, Davis, P30 AG010129; University of California, Irvine, P50 AG016573, P50, P50 AG016575, P50 AG016576, P50 AG016577; University of California, Los Angeles, P50 AG016570; University of California, San Diego, P50 AG005131; University of California, San Francisco, P50 AG023501, P01 AG019724; University of Kentucky, P30 AG028383, AG05144; University of Michigan, P50 AG008671; University of Pennsylvania, P30 AG010124; University of Pittsburgh, P50 AG005133, AG030653; University of Southern California, P50 AG005142; University of Texas Southwestern, P30 AG012300; University of Miami, R01 AG027944, AG010491, AG027944, AG021547, AG019757; University of Washington, P50 AG005136; Vanderbilt University, R01 AG019085; and Washington University, P50 AG005681, P01 AG03991. The Kathleen Price Bryan Brain Bank at Duke University Medical Center is funded by NINDS grant # NS39764, NIMH MH60451 and by Glaxo Smith Kline. Genotyping of the TGEN2 cohort was supported by Kronos Science. The TGen series was also funded by NIA grant AG034504 to AJM, The Banner Alzheimer’s Foundation, The Johnnie B. Byrd Sr. Alzheimer’s Institute, the Medical Research Council, and the state of Arizona and also includes samples from the following sites: Newcastle Brain Tissue Resource (funding via the Medical Research Council, local NHS trusts and Newcastle University), MRC London Brain Bank for Neurodegenerative Diseases (funding via the Medical Research Council),South West Dementia Brain Bank (funding via numerous sources including the Higher Education Funding Council for England (HEFCE), Alzheimer’s Research Trust (ART), BRACE as well as North Bristol NHS Trust Research and Innovation Department and DeNDRoN), The Netherlands Brain Bank (funding via numerous sources including Stichting MS Research, Brain Net Europe, Hersenstichting Nederland Breinbrekend Werk, International Parkinson Fonds, Internationale Stiching Alzheimer Onderzoek), Institut de Neuropatologia, Servei Anatomia Patologica, Universitat de Barcelona. ADNI Funding for ADNI is through the Northern California Institute for Research and Education by grants from Abbott, AstraZeneca AB, Bayer Schering Pharma AG, Bristol-Myers Squibb, Eisai Global Clinical Development, Elan Corporation, Genentech, GE Healthcare, GlaxoSmithKline, Innogenetics, Johnson and Johnson, Eli Lilly and Co., Medpace, Inc., Merck and Co., Inc., Novartis AG, Pfizer Inc, F. Hoffman-La Roche, Schering-Plough, Synarc, Inc., Alzheimer’s Association, Alzheimer’s Drug Discovery Foundation, the Dana Foundation, and by the National Institute of Biomedical Imaging and Bioengineering and NIA grants U01 AG024904, RC2 AG036535, K01 AG030514. We thank Drs. D. Stephen Snyder and Marilyn Miller from NIA who are *ex-officio* ADGC members. Support was also from the Alzheimer’s Association (LAF, IIRG-08-89720; MP-V, IIRG-05-14147) and the US Department of Veterans Affairs Administration, Office of Research and Development, Biomedical Laboratory Research Program. P.S.G.-H. is supported by Wellcome Trust, Howard Hughes Medical Institute, and the Canadian Institute of Health Research. Data for this study were prepared, archived, and distributed by the National Institute on Aging Alzheimer’s Disease Data Storage Site (NIAGADS) at the University of Pennsylvania (U24-AG041689-01), funded by the National Institute on Aging.

**NACC:** The NACC database is funded by NIA/NIH Grant U01 AG016976. NACC data are contributed by the NIAfunded ADCs: P30 AG019610 (PI Eric Reiman, MD), P30 AG013846 (PI Neil Kowall, MD), P50 AG008702 (PI Scott Small, MD), P50 AG025688 (PI Allan Levey, MD, PhD), P50 AG047266 (PI Todd Golde, MD, PhD), P30 AG010133 (PI Andrew Saykin, PsyD), P50 AG005146 (PI Marilyn Albert, PhD), P50 AG005134 (PI Bradley Hyman, MD, PhD), P50 AG0 16574 (PI Ronald Petersen, MD, PhD), P50 AG005138 (PI Mary Sano, PhD), P30 AG008051 (PI Steven Ferris, PhD), P30 AG013854 (PI M. Marsel Mesulam, MD), P30 AG008017 (PI Jeffrey Kaye, MD), P30 AG010161 (PI David Bennett, MD), P50 AG047366 (PI Victor Henderson, MD, MS), P30 AG010129 (PI Charles DeCarli, MD), P50 AG016573 (PI Frank LaFerla, PhD), P50 AG016570 (PI Marie-Francoise Chesselet, MD, PhD), P50 AG005131 (PI Douglas Galasko, MD), P50 AG023501 (PI Bruce Miller, MD), P30 AG035982 (PI Russell Swerdlow, MD), P30 AG028383 (PI Linda Van Eldik, PhD), P30 AG010124 (PI John Trojanowski, MD, PhD), P50 AG005133 (PI Oscar Lopez, MD), P50 AG005142 (PI Helena Chui, MD), P30 AG012300 (PI Roger Rosenberg, MD), P50 AG005136 (PI Thomas Montine, MD, PhD), P50 AG033514 (PI Sanjay Asthana, MD, FRCP), P50 AG005681 (PI John Morris, MD), and P50 AG047270 (PI Stephen Strittmatter, MD, PhD).

